# Predation risk differentially affects aphid morphotypes: impacts on prey behavior, fecundity and transgenerational dispersal morphology

**DOI:** 10.1101/2021.08.30.458248

**Authors:** Sara L. Hermann, Sydney A. Bird, Danielle R. Ellis, Douglas A. Landis

**Affiliations:** Center for Chemical Ecology, Department of Entomology, The Pennsylvania State University, University Park USA; Department of Entomology, Michigan State University, East Lansing USA; Program in Ecology, Evolutionary Biology and Behavior, Michigan State University

**Keywords:** Non-consumptive effects, non-lethal effects, transgenerational effects, trait-mediated interactions

## Abstract

To avoid predation, prey initiate anti-predator defenses such as altered behavior, physiology and/or morphology. Prey trait changes in response to perceived predation risk can influence several aspects of prey biology that collectively contribute to individual success and thus population growth. However, studies often focus on single trait changes in a discrete life stage or morphotype. We assessed how predation risk by *Harmonia axyridis* affects several important traits in the aphid, *Myzus persicae*: host plant preference, fecundity and investment in dispersal. Importantly, we examined whether these traits changed in a similar way between winged (alate) and wingless (apterous) adult aphid morphotypes, which differ in morphology, but also in life-history characteristics important for reproduction and dispersal. Host plant preference was influenced by the presence of *H.axyridis* odors in choice tests; wingless aphids were deterred by the odor of plants with *H.axyridis* whereas winged aphids preferred plants with *H.axyridis* present. Wingless aphids reared in the presence of ladybeetle cues produced fewer offspring in the short-term, but significantly more when reared with exposure to predator cues for multiple generations. However, winged aphid fecundity was unaffected by *H.axyridis* cues. Lastly, transgenerational plasticity was demonstrated in response to predation risk via increased formation of winged aphid morphotypes in the offspring of predator cue-exposed wingless mothers. Importantly, we found that responses to risk differ across aphid polyphenism and that plasticity in aphid morphology occurs in response to predation risk. Together our results highlight the importance of considering how predation risk affects multiple life stages and morphotypes.

## INTRODUCTION

Among the most important interspecific interactions in ecology is the ongoing battle between predators and prey. The complexity of these interactions has been emphasized in the past several decades as research has demonstrated the importance of non-consumptive predator effects – that is, the overall impact of anti-predator decision making on prey survival and performance (Lima 1998). Once the threat of predation is detected, prey can initiate changes in behavior and physiology that help avoid attack or allow them to be less conspicuous to predators over time (Lima and Dill 1990, Stankowich and Blumstein 2005). In addition to changes in behavior and physiology, prey can also exhibit plasticity in morphological diversity, often inducing defenses that limit predator success (Agrawal et al. 1999). While ecological theory increasingly includes the impact of non-consumptive effects in attempts to explain the abundance and distribution of animals across taxa and environments (Peacor et al. 2013), there is still much to explore concerning the influence of predation risk on prey trait plasticity. Consideration of how multiple traits might change in prey organisms is crucial to understanding the impact of predation risk on overall fitness (DeWitt and Langerhans 2003, Preisser and Bolnick 2008). Specifically, there remains a dearth of research that considers predator-induced phenotypic plasticity across multiple traits while considering the life stage or morphology of prey.

The role of predation risk on anti-predator decision making by prey and resulting non-consumptive effects have been demonstrated primarily in aquatic insect and fish systems as well as in several terrestrial mammalian systems (Preisser et al. 2005), leaving much to be explored in terrestrial insects (Hermann and Landis 2017). Among insects, aphids represent a unique group with a complex life-history. While many aphid species lay eggs as needed for overwintering, their dominant form of reproduction is asexual live birth to nymphs. Furthermore, aphids exhibit a form of polyphenism where aphid mothers can generate two distinct adult morphotypes which vary in life history strategy – one of which is a winged morph (alate) upon adulthood, primarily for dispersal and the other wingless morph (aptera) is a more sedentary and primarily reproductive morphotype (Blackman and Eastop 2000, Braendle et al. 2006). There is evidence that apterous aphids alter reproduction and host preferences in response to predators and their cues (Dixon and Agrawala 1999; Fill et al, 2012; Kaplan and Thaler; Nincovik 2013). In general, aptera tend to produce more offspring than their winged counterparts since there are significant reproductive trade-offs associated with the production of wings in alates (Johnson 1963, Groeters and Dingle 1989). Since there are physiological differences between morphotypes, we might predict that the induction of wings can lead to variation in other phenotypes as well as energetic tradeoffs required for wing formation. In addition, because alate aphids are responsible for long-distance dispersal and colonization, if we wish to appropriately model population dynamics of these pest species, it is crucial to understand how alate aphids repond to risk.

The formation of alates in aphid populations is generally considered a response to stressors (plant quality, overcrowding, or pathogens) that allows for dispersal from adverse conditions (Müller et al. 2001, Kunert and Weisser 2005, Hatano et al. 2012; Mehrparvar et al. 2013). There are examples of predator-induced wing formation in aphids, though most studies have focused on a single species of aphid, *Acyrthosiphum pisum* Harris, in direct contact with its natural enemies (Dixon and Agarwala 1999, Weisser et al. 1999, Mondor et al. 2005, Kunert and Weisser 2005, Kaplan and Thaler 2012, Purandare et al. 2014, Kersch-Becker and Thaler 2015, But see: Mehrparvar et al. 2013 for a non-pea aphid example). Interestingly, while it is clear that aphid mothers can induce transgenerational plasticity in response to physical contact with predators, experiments examining the effects of predators on aphid traits have focused exclusively on the apterous morph to date. Transgenerational effects of predation risk have been examined largely in vertebrate systems but have also been demonstrated to influence grasshopper locomotion in an insect system (Hawlena et al. 2011). It remains unclear if the alate morphotypes, which exhibit a dramatically different life history strategy and disperse to generate new populations across the landscape, also exhibit plasticity in their behaviors and reproduction similar to that of the apterous morphotype. The relative importance of response to predation risk and predator cues could vary between these two life histories with potentially less impacts on aphids invested in dispersal and a stronger impact on aphids which are more sedentary and invested in reproduction. To our knowledge, there is no comparison of the impact of predators or predator cues across this polyphenism in aphids. In order to understand the full impact of predation risk on aphids, it is crucial to understand how life history strategy of the prey might affect responses to predation risk.

Our objective was to understand whether predation risk differentially influences phenotypically plastic traits in different morphotypes of the same species. To that end, we assessed how alate and apterous aphids respond to predator cues in several biologically relevant traits (behavior, reproduction and morphology) in both alate and apterous aphid morphs. We utilized green peach aphid (*Myzus persicae* Sulzer) prey and multi-colored Asian ladybeetle (*Harmonia axyridis* Pallas) predators to first ask if the presence of these predators or predator cues on plants alters host plant preference and if the responses differed between alate and apterous morphs. Then, we evaluated the impact of predator cues on aphid fecundity in both morphs, both in the short-term and across multiple generations. Lastly, we asked if the presence of predator odor cues from *Harmonia axyridis* would influence aphid investment in dispersal by inducing the production of alate morphs.

## METHODS

### Plants and Insects

A colony of *M. persicae* was maintained on *Brassica oleracea* Linnaeus (cv. Georgia collard greens) in a climate-controlled insectary (22 C; 16:8 L:D photoperiod). Collard host plants in colony cages were watered weekly and replaced periodically to avoid aphid crowding. Cages contained all ages of aphids and alate or apterous adults were collected from these cages as needed for experiments. To control for the age of aphids used in experiments, groups of adults were placed on a fresh host plant and left to reproduce for 24 hours. We would then remove the adults and rear the immature aphids to adulthood for use in experiments.

A colony of *H. axyridis* was established from larval and adult beetles that were field collected in Ingham County, Michigan. All stages of *H. axyridis* were fed a mixture of corn leaf aphids (*Rhopalosiphum maidis* Fitch) and bird cherry-oat aphids (*Rhopalosiphum padi* Linnaeus) which were reared on barley (*Hordeum vulgare* Linnaeus) plants in 10 cm diameter pots. Colony cages were stored in a climate-controlled growth chamber (25 C; 16:8 L:D photoperiod). Only adult male and female *H. axyridis* were used in experiments.

*M. persicae* colonies (described above) and *B. oleracea* plants were used in experiments. Plants were grown from seed (Burpee, product #52159A) in Promix potting soil (Premier

Horticulture Inc., Quakertown, PA, USA). Germinating seeds were placed in a climate-controlled greenhouse (25 C; 16:8 L:D) and watered daily. Once plants were established, stems were thinned to one plant per cell in a 100-cell plug tray and fertilized once weekly (20-20-20, Peters Professional Water-Soluble Fertilizer, Brantford, Ontario). Once plants were 2-3 weeks old and seedlings had developed true leaves, they were transferred from plug trays to 10 cm round pots where they remained until use in experiments at 4-6 weeks old.

### Aphid Host Cue Preference in the Presence of Predator Cues

Two-arm olfactometer experiments were designed to determine the effects of ladybeetle volatile odor cues on the behavior of the prey insect, *M. persicae* (for a detailed diagram, see **Figure 1A**). All experiments were conducted in a climate and light controlled walk-in growth chamber (25 C, 16:8 L:D photoperiod). Odor sources were placed in 35 cm tall, 615 cm wide dome-shaped glass arenas (ARS, Gainseville, Florida) set atop teflon guillotines and connected to 1.0 L/min, charcoal filtered, and humidified air flow. Guillotines were placed around the stem of the plant, sitting on the rim of the pot, allowing the foliage of the plant to enter the glass arena but excluding the pot, soil and base of the plant. Two separate odor source arenas were set-up in tandem, one for control and one for an odor treatment, 16 h prior to experimentation to allow for plant and insect acclimatization and volatile cue build-up. Control and treatment arenas were then connected via teflon tubing with each odor source supplementing airflow to an individual arm at the end of a y-shaped olfactometer (y-tube). The olfactometer consisted of an 11 cm long glass tube that branched into two 7.5 cm arms. The internal diameter of the tube and arms was 1.5 cm. In this way, each arm of the “Y” consisted of a distinct odor source that flowed down towards the base of the “Y” where insects were released and left to make a choice. Prior to each assay, we collected adult aphids from the colony and confirmed their life stage by detecting the presence of the anal plate under a dissecting microscope. For each experimental replicate, a single adult aphid was selected randomly and placed at the open end of the olfactometer with a fine-tipped paintbrush. Aphid movement towards either the treatment or control arm was observed for a maximum of fifteen minutes. Overall responses were high; we only recorded 4 apterous aphids and 8 alate aphids that did not make a choice after the allotted time. A choice was recorded when the herbivore moved at least halfway up one of the branched arms of the olfactometer. One replicate was conducted per individual aphid. Following each replicate, the y-shaped olfactometer was washed with both acetone and hexane and left to dry to ensure that aphids were not influenced by the movement of their conspecifics in the glassware during previous replicates. Odor sources were changed out after every 10 aphid replicates. In addition, the treatment and control tubes were switched from right to left arm of olfactometer prior to each trial in order to reduce positional bias. All trials were conducted between 09:00 and 13:00 hours.

**Figure 1.**
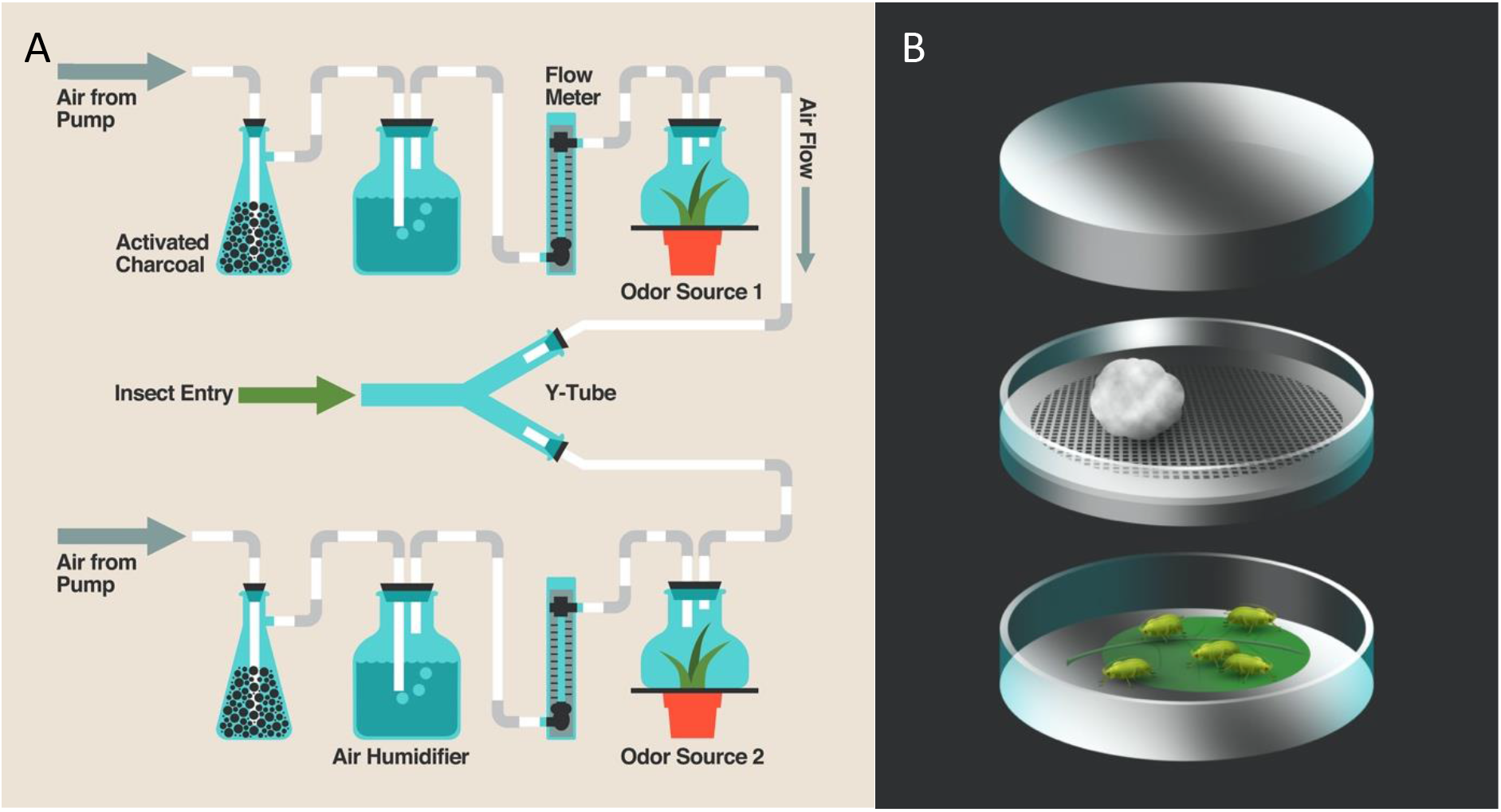
Experimental apparatus used to detect aphid response to predator cues. (A) Schematic of the Y-tube olfactometer set-up. Air flows first through charcoal filter, is regulated using a flow-meter, then humidified using a flask filled with distilled water and finally pumped into the glass chamber which contained odor treatments. Air is then pumped from the odor treatment chamber directly into one of the arms of the “Y”. Aphids were placed individually at the base of the “Y” and monitored for their first choice into one of the arms of the olfactometer. (B) Modified petri dish used to examine aphid nymph production and alate formation in response to predator cues or predator-free controls. Right: the separated portions of the petri dish included the bottom, which contained a moistened filter paper and a leaf disc where aphids were placed; the center which contained two modified petri dish lids that held a mesh barrier between the aphid prey and the treatments; the top, this is the portion that contained the treatments which were either 1) a moistened cotton ball (control) or 2) a moistened cotton ball with two *H. axyridis* adults (predator). (Diagrams courtesy of Nick Sloff, Department of Entomology, The Pennsylvania State University).

#### Odor treatments

The odor sources for all y-tube assays used the same basic arena set-up which consisted of a single collard plant and a moistened cotton ball placed inside the glass chamber (described above). This served as the control odor source. We also created two predator odor treatments, ‘predator + plant’ treatment and ‘predator pre-treatment’. For both we used the same basic set-up and added five male and five female ladybeetles for 16 h prior to experimentation. Predators remained on the plant during the y-tube assay for the predator + plant treatment but were removed just prior to the assays for the predator pre-treatment odor source. In this way we were able to control for potential indirect effects of predators on plant odors. Y-tube assays were run with a control odor source and one of the two predator treatments for both apterous (N=50) adult and alate (N=35) *M. persicae*.

### Aphid Performance in Response to Predator Cues in Petri Dish Arenas

We examined whether *M. persicae* would alter the number of nymphs they produce in the presence of predator cues using a modified petri dish arena. In this experimental arena, we physically separated aphid prey from ladybeetle predators while allowing volatile odors and visual cues of these predators to be experienced by the developing aphids. Petri dish arenas were made by cutting a 7 cm diameter hole in the larger half of two petri dishes. The lids were placed top to top enclosing a mesh screen and fixed together with hot melt glue (**Figure 1B**). A freshly excised collard leaf disc (60 mm diameter) was placed directly atop moistened filter paper (Whatman 90 mm circles) cut to fit the bottom portion of the petri dish arena. One of two treatments was placed in the top portion of the petri dish experimental arenas: 1) predator-free control treatment with a single moistened cotton ball or 2) predator treatment with two *H. axyridis* adults and a single moistened cotton ball. For each experimental replicate, a single apterous or alate aphid adult was left to reproduce for 3 d. At the end of the experiment, we counted the number of nymphs produced. For apterous aphids, 51 replicates of each treatment were performed; for alates there were n = 59 control and n = 58 predator cue replicates.

### Alate Formation in Response to Predator Cues in Petri Dish Arenas

We used the modified petri dishes (described above) to examine if predation risk affects aphid physiology. Here, we exposed aphids to predator cues for 3 d and monitored for induction of alate morphs. One of two treatments was placed in the top portion of the petri dish experimental arenas as described above. In each arena, five adult apterous aphids were randomly selected from the stock aphid colony and gently placed on the leaf disc with a fine-tipped paint brush. Aphids were then exposed to either the control or predator treatment continuously for 3 d. after which the total number of aphids that developed wings in each treatment were counted. There were 20 replicates for each treatment.

### Influence of Predator Cues on Aphid Fecundity and Alate Formation on Intact Plants

We also examined the impact of predator cues on nymph production and alate formation on intact plants over a longer duration of time. For this experiment, we utilized 4 w old collard plants grown in 5.08 cm diameter round pots. Potted plants were placed inside 710 ml cylindrical glass ball jars (Ball, item # 1033893) on top of one sheet of filter paper (Whatman 90mm circles). For each replicate, seven apterous adult aphids were chosen randomly from the stock colony and placed on the plants inside the jars. In each ball jar, we placed a mesh barrier between the plant and the lid of the jar, where treatments were placed. A mesh barrier was fashioned approximately 3 cm above the top of the plant by inserting a plastic acetate ring that fit snugly in the top portion of the ball jar arena. On the top and bottom of the acetate ring, mesh was used to allow for airflow and exposure to treatments while also prohibitting aphid or ladybeetle movement out of the arena.

Three treatments were established: 1) a control treatment with only moistened cotton in the mesh barrier (n = 17), 2) a lethal predator treatment with one male and one female ladybeetle contained within the arena along with the aphids and the host plant (n = 16), and 3) the predator cue “risk” treatment in which one male and one female ladybeetle were separated from aphids by the mesh barrier (n = 18). Jars were sealed with metal ring lids that secured the mesh barrier onto the top of the jar. Jars were placed in a climate-controlled growth chamber as described above for the duration of the experiment. After 7 d, aphids in each jar were counted and the jars were then returned to the growth chamber for an additional 7 d. After the second 7 d period, jars were removed from the growth chamber and plants were removed from jars in order to obtain a total aphid count and assess alate formation over 14 d. Since *M. persicae* in our colony generally complete a full life cycle in 7 d, this trial represents 1-2 generations of aphid production.

### Statistical Analysis

All data were analyzed using JMP (JMP Pro, Version 12. SAS Institute Inc., Cary, NC, 1989-2007). The number of *M. persicae* entering the control versus treatment arm in the y-tube olfactometer bioassays were compared with chi-square tests. The null hypothesis was equal entrance by aphids into both arms of the olfactometer. We used Fishers exact test to compare the proportion of alates present in the predator treatment to that of the control treatment in both the short-term petri dish assay and the full-plant assay. Here, we predicted the number of alates would differ between treatments, with more produced in response to odor cues of their predators. Data obtained from the remaining experiments were not normally distributed, and we were unable to normalize these data through square root or log transformation, precluding parametric tests. Therefore, we used the non-parametric Wilcoxon signed-rank test to analyze whether nymph production by both alate and apterous *M. persicae* differed from the null hypothesis of equal numbers of offspring between treatments. Finally, our longer-term nymph production and alate formation experiment data were first analyzed to compare the number of aphids across the three treatments using a Kruskal-Wallis one-way analysis of variance. Then, each treatment pair was analyzed using a post-hoc non-parametric Wilcoxon multiple comparisons test.

## RESULTS

### Aphid Host Cue Preference in the Presence of Predator Cues

When presented with a choice between a predator-free odor source or an odor source that included *H. axyridis* predators, 66% of adult apterous *M. persicae* preferred the arm with predator-free control plants (*χ* ^2^ (n = 50) = 5.12, p = 0.024, **Figure 2**). However, when the physical predators were removed from the odor source arena prior to bioassays, the adult apterous aphids no longer preferred predator-free control plants (*χ* ^2^ (n = 31)= 3, p = 0.083, **Figure 2**). In contrast, 71% of the alate *M. persicae* preferred to move towards plants with predators present compared to the predator-free odor source (*χ* ^2^ (n = 35) = 7.53, p = 0.006, **Figure 2**), but only when the physical predators were in the odor source arena. When predators were removed from the odor source arena prior to bioassays, equal preference between the olfactometer arms was observed (*χ* ^2^ (n = 31) = 0.037, p = 0.847, **Figure 2**).

**Figure 2.**
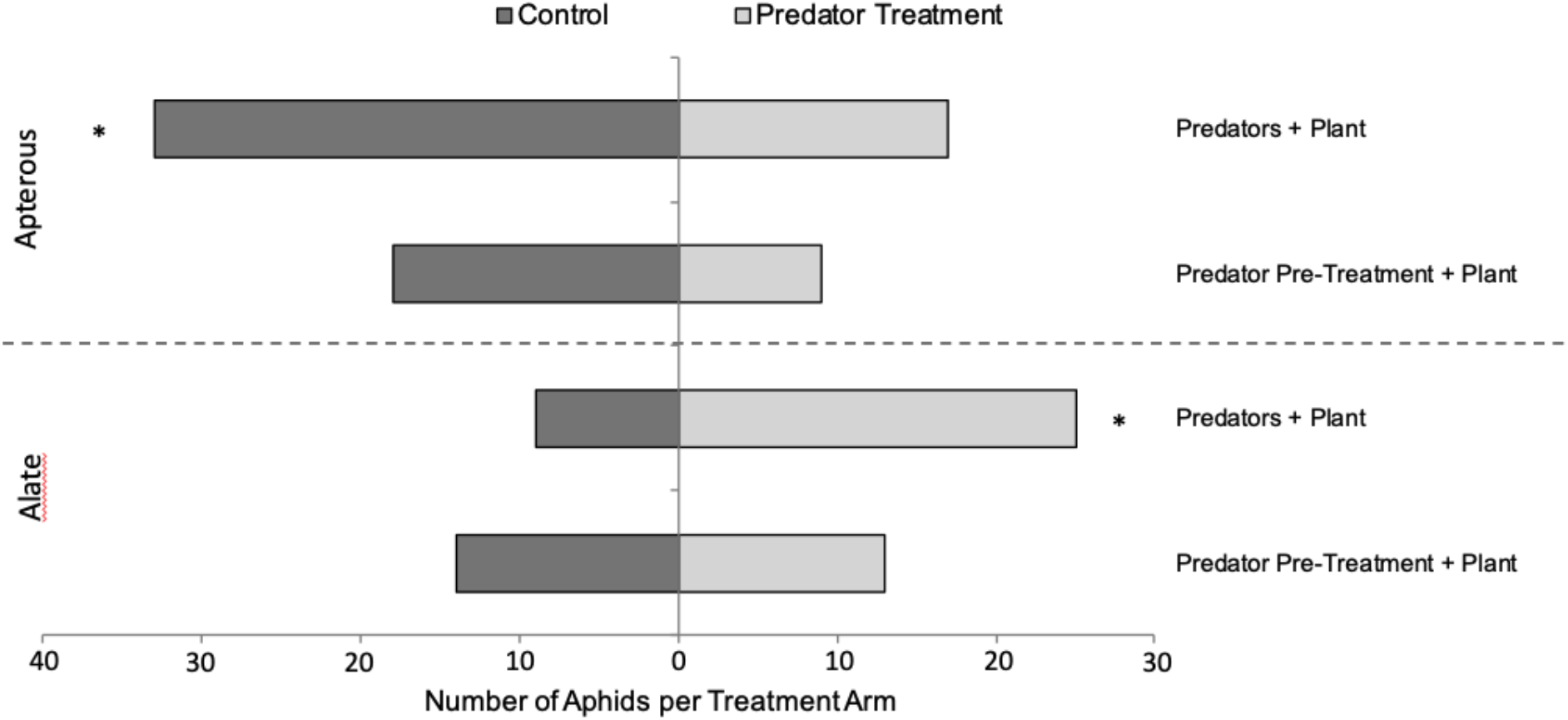
Responses of adult apterous and alate *M. persicae* to odor sources in a two-choice y-tube olfactometer (top). Treatment plants were pre-exposed to 10 *H. axyridis* adult predators for 16 h and control plants were predator-free (* indicates significance at p < 0.05 following chi-square test of goodness of fit).

### Aphid Performance in Response to Predator Cues in Petri Dish Arenas

Adult apterous *M. persicae* exposed to predator cues from *H. axyridis* in a petri dish arena had a 23% reduction in the overall number of nymphs produced over 3 d compared to reproducing adult aphids in control petri dishes where predator cues were absent (Z = -4.08, p < 0.0001, **Figure 3A**). In contrast, when adult alate *M. persicae* were left to reproduce in the presence of predator cues there was no discernible effect on nymph production compared to predator-free controls (Z = -0.46, p = 0.65, **Figure 3B**).

**Figure 3.**
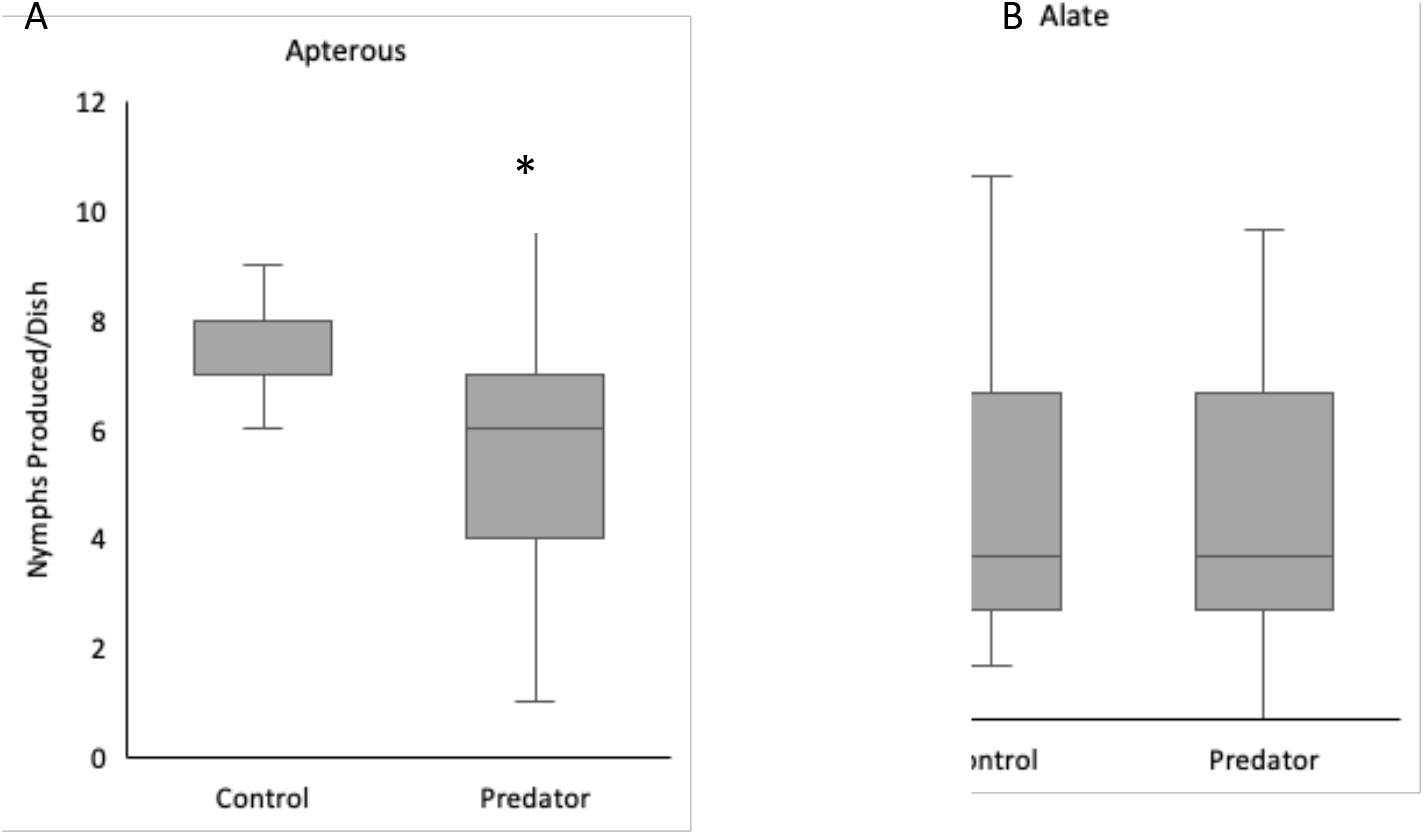
Nymph production by single *M. persicae* (A) apterous morphs or (B) alate morphs in a petri dish arena. Aphids were exposed to either a predator-free control or a predator treatment consisting of two *H. axyridis* ladybeetle predators for three consecutive days (* indicates significance at p < 0.05 as indicated by the non-parametric Wilcoxon signed-rank test).

### Alate Formation in Response to Predator Cues in Petri Dish Arenas

To investigate the potential for predation risk to induce wing formation, we exposed aphids to predator cues by physically separating the aphids on leaf discs from ladybeetle predators in a petri dish. In this experiment, the number of individuals that produced wings after 3 d in petri dishes differed between the predator cue treatment and the predator-free control, with a five-fold increase in alate production in the predator cue treatment (p = 0.039; Control: 3, Predator Risk: 15). Overall, 3% of aphids in the control treatment developed into alate adults by 3 d, whereas 15% of aphids formed wings in the predation risk treatment dishes that left aphids exposed to predator cues.

### Influence of Predator Cues on Aphid Fecundity and Alate Formation on Intact Plants

Nymph production differed significantly among treatments (χ^2^ = 32.87, p < 0.0001, **Figure 4**). Pairwise comparisons of the different treatments show that the risk treatment yielded significantly more nymphs than the control and lethal treatments (Z = 3.219, p = 0.0013; Z = 4.903, p < 0.0001, respectively) while lethal treatment had the fewest aphids after 14 d. Alate formation was significantly increased in the risk treatment (n = 12 individuals) compared to both the control and lethal treatment where no alates were found during the entire experiment (G = 16.636, p < 0.0001).

**Figure 4.**
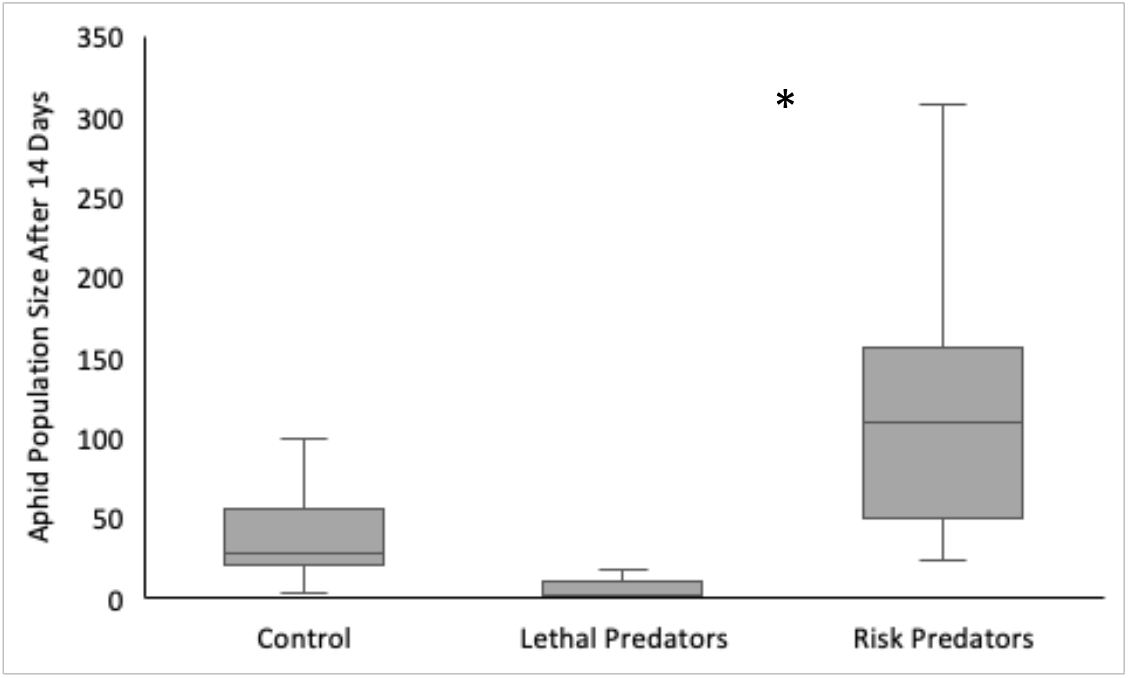
Nymph production by seven *M. persicae* apterous aphids exposed to either a predator-free control, two *H. axyridis* ladybeetle predators (lethal), or two *H. axyridis* ladybeetle predators confined in a mesh barrier (risk) for 14 consecutive days (* indicates significance at p < 0.05 following a non-parametric Kruskal-Wallis, one-way analysis of variance).

## DISCUSSION

Our results demonstrate that *M. persicae* exhibits plasticity in several important traits when exposed to *H. axyridis* predator cues. Importantly, we found that predation risk has differential effects on alate versus apterous aphids which vary in both their morphology and life history. We observed behavioral preferences in aphid orientation to host odor cues when given the choice between predator-free host odors and host plants with predators present. Interestingly, while apterous morphs avoided predators on plants by choosing to walk towards predator-free controls, alate aphids preferentially move towards plants that harbored predators. In the presence of predator cues, apterous aphid fecundity was altered by initially reducing nymph production (3 d) and subsequently increasing nymph production when in the presence of predator cues for a longer period (14 d) representing multiple generations. However, alate aphid fecundity over 3d did not differ in the presence of predatory cues compared to controls and we thus we did not explore alate fecundity in the long-term assay. Lastly, we found increased investment in the formation of dispersal morphs in the offspring of aphids in the presence of predator cues, representing transgenerational impacts of risk exposure. Together, these results show that aphid prey are capable of using predator cues to identify risk and respond by altering behavior, fecundity and morphology, but that anti-predator changes in traits differ between the two aphid morphotypes highlighting that life history strategy might influence response to risk.

Apterous aphid adults avoided plants that harbored predators and strongly preferred predator-free control plants. There was also a trend for these aphids to avoid plants that had previously harboured predators. Because apterous aphids lack wings and thus the ability to disperse by flight, preference for a predator-free plant would be adaptive and provide offspring an environment that is free of enemies (Lee et al. 2011, Wasserberg et al. 2013, Sendoya et al. 2015, Hermann and Thaler 2018). Aphid movement between plants by apterous aphids can be an important dispersal strategy in some species of aphids (Losey and Denno 1998, Kersch-Becker and Thaler 2015). To understand if the preference we found in the y-tube olfactometer would allow for increased dispersal away from predation risk, future experiments where aphids can move freely between risky and control plants will be necessary. In addition, apterous adults reduced their production of nymphs in the presence of close-range predator cues over 3 d. This result followed our initial expectation that investment in offspring would be reduced by detection of predation risk. While giving live-birth, aphids are likely less able to move and defend themselves and thus, either avoiding plants that contain predators or reducing apparency by altering behavior would be a strategy for predator evasion. One caveat regarding this assay was that it was performed in small arenas and thus cues were very concentrated and spatially confined, which may not be representative of this system in nature. Interestingly, when we scaled this experiment up to provide prey with a full plant, rather than a leaf-disc, and exposed them to the same predator cues for a longer period of time (representing 1-2 generations), apterous adults produced significantly more nymphs compared to the no-predator control. Because the adult aphids in this experiment were unable to disperse by walking to a predator-free plant, perhaps here their strategy shifts to one of bet-hedging with long-term exposure to predator cues (Grégoir et al. 2018). In this case, the more offspring produced by individual adults might allow for the population to succeed, even in the face of predation risk. Increased production of offspring under predation pressure has been found in at least one other aphid system. Potato aphids (*Macrosiphum euphorbiae* Thomas) were exposed to convergent lady beetle predators (*Hippodamia convergens* Guérin-Méneville) that were rendered non-lethal through mouthpart manipulation, significantly higher numbers of nymphs were produced by the aphids (Kersch-Becker and Thaler 2015). It has also been shown that Colorado potato beetle (*Leptinotarsa decemlineata* Say) response to stink bug (*Podisus maculiventris*) predator cues can vary across time in a field setting, with a decrease in altered prey feeding behavior over the season (Aflitto and Thaler, 2020). Therefore, it is possible that habituation to predator odor cues, especially in the absence of aphid alarm cues, relaxed the impact of risk on aphid reprodiction. As the field of non-consumptive effects of predators on prey continues to expand, it will be important to better understand the factors that influence the directionality of prey trait changes in response to risk.

In contrast to our findings with aptera, alate aphids were attracted to host plants with predators in our y-tube choice experiments, which was contrary to our predictions that all prey morphotypes would avoid plants with predator cues associated with them. In another system, *L.decemlineata* colonization of potato fields was not affected by the presence of predators *P.maculiventris,* yet subsequent behaviors such as feeding were altered once prey were established on plants (Hermann and Thaler 2018). While an attraction to host plants harboring predators might not intuitively be adaptive, it is possible that alates are better equipped to avoid predators on plants due to the presence of wings. Conversely, it is also possible that alate attraction to these plants is maladaptive and a result of lady beetles actively attracting prey as a strategy to obtain a food source. In our study, we also measured the fecundity of alate aphids in response to predator cues but there was no difference in nymph production compared to predator-free controls. Since alate aphids are produced in response to a variety of stressors in order to facilitate dispersal and re-colonization of aphid colonies across the landscape, it is also possible that this life stage is less likely to respond to risk overall. In addition, the presence of wings enhances the mobility and potential for escape from predators which could also influence propensity to induce anti-predator changes in reproduction. In future studies, it will be necessary to monitor the outcome of alate colonists on plants that contain predators. Further, the attraction to host plants by alates is no longer significant when predators are removed prior to experiments, suggesting that physical presence of predators is necessary for the attraction to occur. Again, to gain insight on this result, work must be done to elucidate the adaptive potential of choosing a plant where predators are actively foraging.

Aphid investment in producing a higher proportion of dispersal morphs in response to various stressors (plant quality, crowding, alarm cues, natural enemies) has been previously demonstrated and modelled (Dixon and Agarwala 1999, Weisser et al. 1999, Kunert and Weisser 2003, Mondor et al. 2005, Kaplan and Thaler 2012, Kersch-Becker and Thaler 2015; Poethke et al. 2010). In our study, we found that alate formation was higher in the presence of predator cues compared to controls in both our petri-dish and full plant assays. This result is demonstrated in our full plant experiment because alates were only found in the risk predator treatment that provided predator cues. There were no alates found in control treatments or treatments with lethal predators. In this experiment, aphid abundance was also highest in the risk treatment and since crowding can lead to increased alate formation (Purandare et al. 2014), the influence of density-dependent alate formation cannot be ruled out. Wing formation could be a result of crowding stress, predator cues or the combination of crowding and predator cues in our experiment. However, previous work has shown that aphids increase wing production in the presence of predators, but only when their antennae are intact (Kunert and Weisser 2005), suggesting that volatile chemical cues are likely responsible for morph induction. Our work in the short term assay highlights that, without crowding, alate formation is induced. Future studies should aim at disentagling the impact of crowding from predator cues in alate formation and dispersal behavior.

Our study adds to a growing body of literature demonstrating that predator cues are a factor in prey detection of predation risk and that detection can lead to varied responses in different morphotypes of the same prey animal. In addition, we show that several prey traits are influenced by predator odor cues, all of which are important for the success of individual aphids and could scale to interfere with the success of the population. Our study further suggests the important role of predator chemical cues in predation risk related non-consumptive effects (Gonthier 2012, Hoefler et al. 2012, Ninkovic et al. 2013, Hermann and Thaler 2014), which not only has direct implications for understanding fundamental insect ecology, but also has practical applications in pest management and conservation efforts (Hermann and Landis 2017) and shows promise in aphid systems (Ingerslew and Finke, 2020). Future work must look at the adaptive potential of these shifts in behavior and physiology to determine if these trait changes ultimately aid in predator avoidance and overall survival or if they are maladaptive and lead to a net negative impact on prey population growth and success.

## Acknowledgements

We thank members of the Landis lab, Jared Ali, Jessica Kansman and two anonymous reviewers for helpful critiques of the manuscript. We also thank Aubrey McElrath, Lane Proctor and Carissa Blackledge for assistance in experimental execution. Support for SLH was provided by a grant from the USDA National Institute for Food and Agriculture grant (2017-68004-26323), MSU Plant Science Fellowship and by the NSF Long-term Ecological Research Program (DEB 1832042) at the Kellogg Biological Station (KBS LTER). Support for DAL was provided by the KBS LTER and by Michigan State University AgBioResearch.

## Statement of Human and Animal Rights

This article does not contain any studies with human participants or animals performed by any of the authors.

